# Paradoxical replay can protect contextual task representations from destructive interference when experience is unbalanced

**DOI:** 10.1101/2024.05.09.593332

**Authors:** Hung-Tu Chen, Matthijs A. A. van der Meer

## Abstract

Experience replay is a powerful mechanism to learn efficiently from limited experience. Despite decades of compelling experimental results, the factors that determine which experiences are selected for replay remain unclear. A particular challenge for current theories is a set of studies reporting “paradoxical” replay of trajectories that are neither experienced recently or about to be chosen. To understand why, we simulated a feedforward neural network with two regimes: rich learning (structured representations tailored to task demands) and lazy learning (unstructured, task-agnostic representations). Rich, but not lazy, representations degraded following unbalanced experience, an effect that could be reversed with replay of non-chosen options. To test if this computational principle can account for the experimental data, we examined the relationship between paradoxical replay and learned task representations in the rat hippocampus. Across two different studies, we found an association between the richness of learned task representations and the paradoxicality of replay. Taken together, these results suggest that paradoxical replay may serve to protect rich representations from the destructive effects of unbalanced experience, and more generally demonstrate a novel interaction between the nature of task representations and the function of replay in artificial and biological systems.

**Significance Statement:** We provide an explicit normative explanation and simulations of the experimentally observed puzzle of “paradoxical” replay of experiences that are neither experienced recently or about to be chosen. We show computationally that such replay can serve to protect certain task representations from the destructive effects of unbalanced experience. We experimentally confirm the main prediction of the theory, that “rich” task representations, measured using representational distance in the rodent hippocampus, show more paradoxical replay compared to “lazy” task representations. Our theory refines the notion of consolidation in complementary learning systems theory in showing that not all task representations benefit equally from interleaving, and provides an example of how the use of replay in artificial neural networks can be optimized.

## Introduction

Replay is a powerful mechanism to maximize what can be learned from limited experience. Decades of experimental work have explored the phenomenon of hippocampal replay -- including its causal importance for various memory-guided behaviors (Aleman-Zapata et al., 2022; Girardeau et al., 2009; Gridchyn et al., 2020; Jadhav et al., 2012; but see Deceuninck & Kloosterman, 2024) and some of the major factors that govern what experience to replay when (Foster & Wilson, 2006; Huelin Gorriz et al., 2023; Liu et al., 2021; Michon et al., 2019). A productive interaction between experimental approaches and theoretical work has resulted in several families of ideas about the content and function of replay in the brain. The oldest idea, “systems consolidation” casts replay as the vehicle for off-line information transfer from the short-term storage of one-shot episodic memories in the hippocampus to longer-term semantic information in the neocortex, solving the computational tradeoff between rapid learning of specific experiences on the one hand and extracting generalities on the other (“complementary learning systems”, Buzsáki, 1989; McClelland et al., 1995; O’Reilly et al., 2014). A different idea, within the framework of reinforcement learning, casts replay as a way to update the values of non-local states and actions that are indirectly affected by a recently experienced prediction error, solving the temporal credit assignment problem offline (Mattar & Daw, 2018; Momennejad et al., 2018; Sagiv et al., 2024; Sutton, 1991). Finally, a common proposal is that during on-line decision making, replay implements a forward planning process in which possible upcoming courses of action are retrieved or simulated (Foster, 2017; Joo & Frank, 2018; Ólafsdóttir et al., 2018; Pfeiffer & Foster, 2013; but see van der Meer & Bendor 2025), an idea which has significant overlap with offline credit assignment (Mattar & Daw 2018, Mattar & Daw 2026).

These three ideas do not only assign different (non-exclusive) functions to replay; they also make specific predictions regarding what replay content would be expected given particular experience. For instance, the systems consolidation idea predicts that replay content is fundamentally retrospective, providing a temporary storage buffer of recent experiences to replay from, optionally prioritized by e.g. salience or prediction error (Moore & Atkeson 1993, Schaul et al. 2016). In contrast, offline credit assignment combines both prospective and retrospective, using past experience to fix those inconsistencies most useful for the task at hand, and the planning idea is fundamentally prospective. However, several experimental studies have shown these accounts to be incomplete, observing replay of trajectories that are neither experienced recently nor about to be chosen (Carey et al., 2019; Gillespie et al., 2021; Gupta et al., 2010; Wimmer et al., 2023). Such “paradoxical replay” is challenging to explain in any of these theories. For instance, in the 2-arm maze of Carey et al. rats systematically replayed the arm of the maze that they did not choose behaviorally, and in the 8-arm maze of GIllespie et al. there are two different paradoxical biases: replay of previously rewarded goal locations even after those locations are no longer behaviorally relevant, and a bias toward other non-recently visited unrewarded locations that increases with time since the last visit.

A largely separate computational literature, rooted in the connectionist networks of the 1980s extending to the deep nets of today, has revealed the benefits of replay for the purposes of speeding up learning in general, and the prevention of forgetting more specifically (Hayes et al., 2021; Mnih et al., 2015; Parisi et al., 2019; van de Ven et al., 2020). In particular, the phenomenon of “catastrophic interference” refers to the destructive effects that weight updates for a new item can have on previously stored weights for old items (French, 1999; McClelland et al., 1995; McCloskey & Cohen, 1989). While many of these studies did not use replay directly, it was noted that the interleaving of different items to be learned, rather than doing batched learning of the same item in a repeated fashion, made learning more resilient to interference between items (Carvalho & Goldstone, 2015; Zhou et al., 2023; but see Flesch et al., 2018; Schlichting et al., 2015). Interestingly, those experimental studies that have found paradoxical replay have in common that experience on the task is both highly stereotyped and highly unbalanced, i.e. animals repeatedly run a specific trajectory in an explicitly non-interleaved manner (Table 1). This pattern suggests the hypothesis that when experience is unbalanced, paradoxical replay serves to prevent repeated learning from that same experience from destroying other memories; however, the extent to which biological replay serves this purpose remains unclear.

**Table 1:** Paradoxical replay tends to occur in those experimental studies that feature highly unbalanced experience. . Each row corresponds to a representative study of rats performing a maze-based task in which the behavioral experience (e.g. maze arm traversals) and replay of multiple alternative spatial trajectories can be compared. **Maze layout / task**: a brief description of the environment and task used in the study. **Experience bias**: the crucial independent variable of interest, capturing the distribution of experience across the available options. *Balanced experience* indicates relatively equal traversal experience (trial count) across options (e.g. approximately equal “left” and “right” choices during T-maze alternation; uniform distribution), whereas *unbalanced experience* refers to repeated experience (traversals) of some options to the exclusion of others (e.g. choosing one arm of a maze over the other). **Replay content**: the dependent variable of interest, describing the relationship between the experimentally observed replay content and behavioral experience. *Paradoxical replay* refers to a dissociation between replay content and experience, e.g. replay content that tracks neither recent past experience or upcoming choice. *Orthodox replay* refers to a correspondence between past or upcoming experience and replay content. Paradoxical replay of non-preferred options is most apparent in tasks where animals sample alternatives unevenly; in contrast, tasks where experience is more balanced tend to yield orthodox replay of current, future, or rewarded trajectories (though reward and task demands can bias content even when trial counts are balanced). *\*Note*: as we discuss in the main text, Gillespie et al. 2021 find two different paradoxical biases in awake replay content: a strong bias toward previously rewarded goal locations and a weaker bias toward other non-recently visited unrewarded locations.

| study | maze layout / task | experience bias | replay content |
| --- | --- | --- | --- |
| **Gupta et al. 2010** | Continuous “Two-T” maze | Alternation (balanced) or Left/Right preferred (unbalanced) | Paradoxical (Left/Right preferred) |
| **Carey et al. 2019** | Discrete-trial single T-maze | Left/Right preferred (unbalanced) | Paradoxical |
| **Gillespie et al. 2021** | Back-and-forth 8-arm radial maze | Single arm rewarded in blocks (unbalanced) | Paradoxical* |
| **Wu et al. 2017** | Linear track with shock zone | unbalanced (animals avoid shock zone) | Paradoxical |
| **Dupret et al. 2010** | Circular open-field “cheeseboard” | balanced | Orthodox |
| **Pfeiffer & Foster 2013** | Square open-field “cheeseboard” | balanced | Orthodox |
| **Shin et al. 2019** | Back-and-forth W-maze alternation | balanced | Orthodox |
| **Michon et al. 2019** | Discrete-trial two-arm maze | balanced | Orthodox |

In this study, we first explore the computational properties of replay for the purpose of preventing memory interference when experience is unbalanced. Using simple 3-layer neural networks that learn a simple contextual discrimination task (context 1: L+R-, context 2: L-R+; Figure 1a, b; analogous to Carey et al. and others) we capitalize on prior work that has shown such networks can solve this task using different strategies with distinct computational tradeoffs. Specifically, “lazy” networks learn quickly but the resulting high-dimensional task representations are less robust to noise in the inputs; “rich” networks learn slower, but the resulting low-dimensional context-driven task representation is more robust to noise (Chizat et al., 2018; Flesch et al., 2022, 2023; Woodworth et al., 2020). We show that unbalanced experience causes memory interference in rich, but not lazy, networks, which can be remedied by paradoxical replay. This representation-specific sensitivity to unbalanced experience enables an experimental test of the hypothesis that biological replay serves to protect against memory interference: we should see more paradoxical replay for those representations that are most vulnerable to interference from unbalanced experience, i.e. rich representations. Using empirical measures of rich vs. lazy representations in the rat hippocampus, we confirm the predicted association between the strength of paradoxical replay and the richness of the learned representation across two different experimental studies. These results provide a link between replay content and the nature of task representations, providing a proof of principle for a potential adaptive function for paradoxical replay.

**Figure 1:**
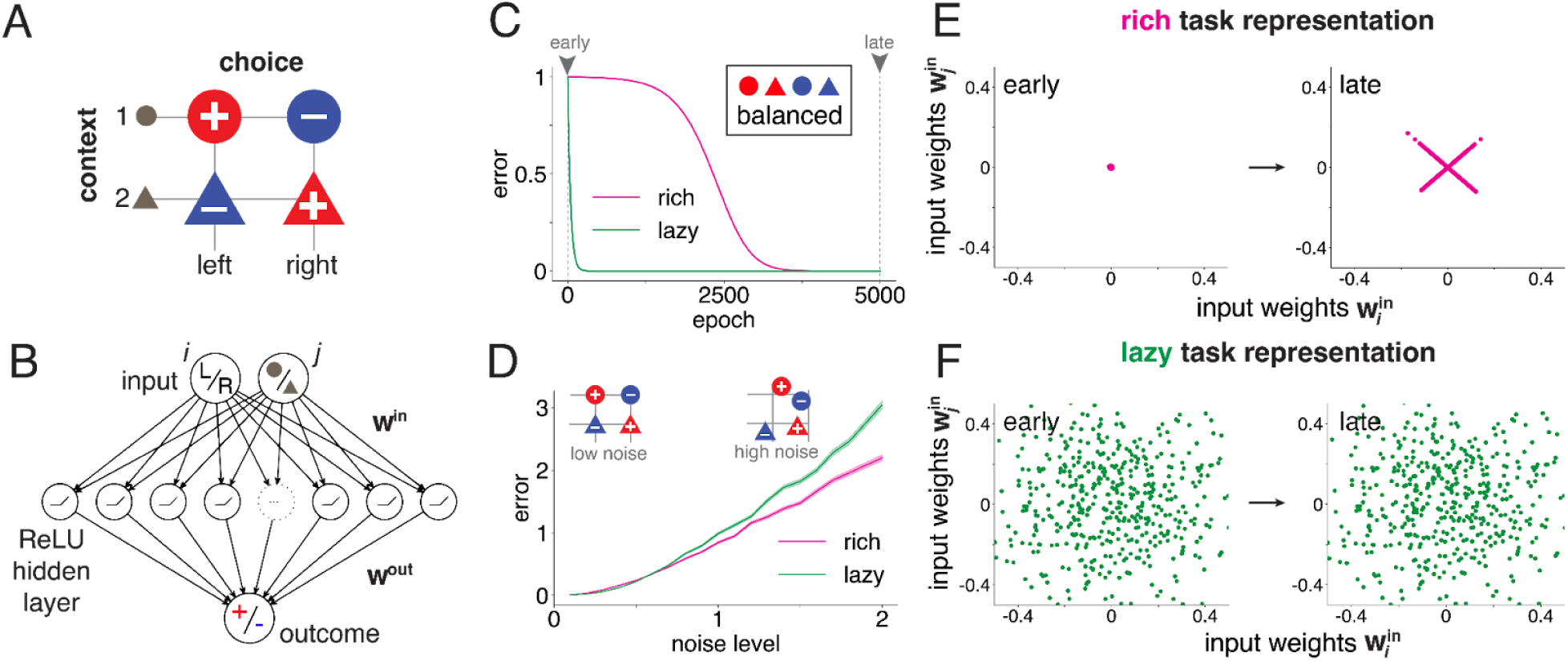
Rich and lazy task representations in a neural network model of a canonical contextual discrimination task. **(A)** Contextual discrimination task schematic: in context 1 (circles), choosing left but not right is rewarded; these contingencies are reversed for context 2 (triangles) **(B)** A feedforward neural network was trained to solve this problem. The network receives inputs of choice *i* (left or right) and context *j* (1 or 2), maps them into a single hidden layer of rectified linear units (ReLU), and ultimately outputs predictions indicating the valence of outcomes (−1, blue; or +1, red). **(C)** Two distinct solutions mediated by different task representations were produced: “rich” networks (magenta) learned slowly from *small* initial weight variance, and “lazy” networks (green) learned quickly from *large* initial weight variance. Training was balanced across all possible context-choice pairs. **(D)** Networks in the rich regime showed greater robustness to input noise compared to those in the lazy regime (shaded area indicates SEM across different runs; insets diagram low-noise and high-noise conditions). **(E)** In rich networks, input-to-hidden weights undergo notable changes, learning contextual, task-related representations specific to the task structure (note the distinctive “x” shape of the weights). **(F)** Conversely, in lazy networks, input-to-hidden weights retain their high-dimensional, task-independent format over the course of training.

## Results

To explore the computational principles governing the interaction between replay, experience, and task representations, we built on the previously described tradeoff between the slow acquisition of “rich” task representations robust to noise on the one hand, and the rapid acquisition of “lazy” but less generalizable task representations on the other (Flesch et al., 2022, 2023). Specifically, we trained 3-layer feedforward neural networks on a canonical contextual discrimination task where the rewarded choice (left or right) depends on context (context 1: L+R-, context 2: L-R+). As expected, rich networks (small initial weight variance, σ = 0.0025) learned more slowly compared to lazy networks (large initial weight variance, σ = 0.25; Figure 1c). Input-to-hidden weights in rich networks captured the task structure in a low-dimensional representation (Figure 1e), which was relatively robust to perturbation in the inputs (Figure 1d). Conversely, lazy networks retained their initial high-dimensional, task-agnostic structure in their hidden layer weights (Figure 1f), with learning primarily occurring in hidden-to-output weights (Figure S1b). This high-dimensional representation in lazy networks made the network more sensitive to input perturbations compared to rich representations (Figure 1d), a distinction that held in both deterministic and noisy training regimes (Figure S1c,d). These results reproduce the previously documented trade-off between robustness to perturbations in the input on the one hand (robust for rich contextual, low-dimensional task representations, fragile for high-dimensional, task-agnostic lazy representations) and speed of learning on the other (slow for rich, fast for lazy representations).

In the above simulations, experience was balanced equally between all possible context and choice pairs. To examine the effects of *unbalanced* training on rich vs. lazy representations, we instead trained these networks with an 80/20 split favoring the rewarded outcome (see Figure S2a and *Methods* for implementation details). Under these unbalanced conditions, rich networks developed biased representations, with both input and output weights favoring positive outcomes (Figure 2b and S2c, left panel; note the difference with Figure 1e) resulting in decreased performance on the task (Figure S2b). Following prior work showing that replay can help prevent catastrophic forgetting (e.g. McClelland et al., 1995; van de Ven et al., 2020) we hypothesized that paradoxical replay could help rebalance representations for both outcomes. To test this, we simulated replays with an overrepresentation of the less-experienced task conditions after biased training (Figure 2c). This paradoxical replay led to unbiased representations for both outcome contingencies (Figure 2d and S2f, left panel), as implied by treating learning from replay as equivalent to learning from real experience. In contrast, lazy networks were insensitive to unbalanced training, and did not require rebalancing using paradoxical replay (Figure 2b, d, right panel). Thus, rich vs. lazy representations were differentially sensitive to unbalanced training, and paradoxical replay can rebalance the representations of a rich learning network in the face of unbalanced training.

**Figure 2:**
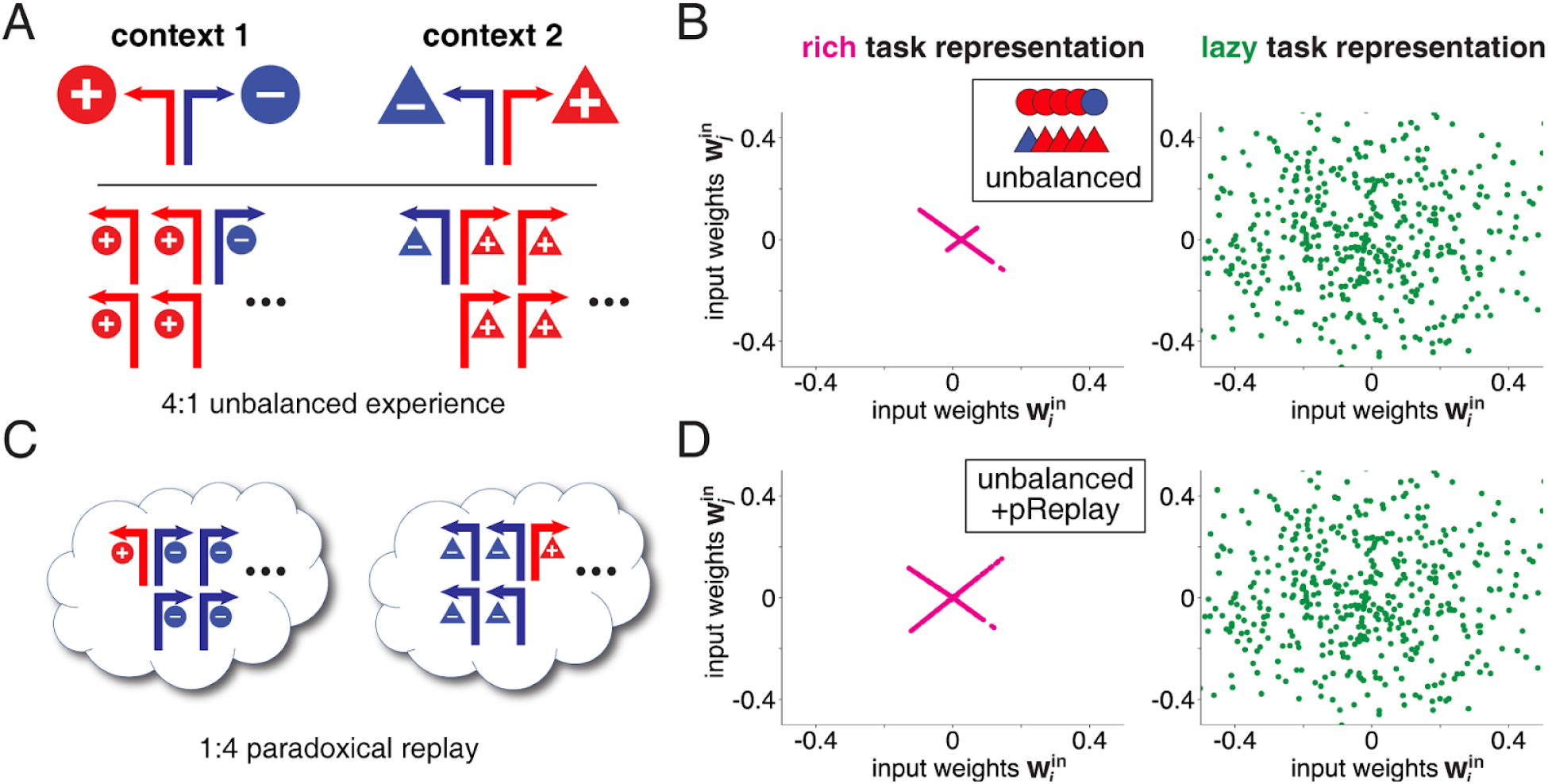
Unbalanced experience degrades rich, but not lazy representations, an effect that can be rescued by paradoxical replay. **(A)** We simulated the effect on task representations given repeated experience of favorable outcomes (left in context 1, right in context 2) using an 80/20 split. **(B)** Biased training on the positive contingency resulted in biased representations of input weights in rich networks (note the asymmetric “x” shape), while lazy networks remained unaffected (compare Figure 1e-f, right panels). **(C)** To illustrate how paradoxical replay might aid networks in retaining a balanced representation of both contingencies, we introduced 1:4 biased replays of the negative (less-experienced) contingency after 4:1 biased training favoring the positive contingency. **(D)** Input weights of rich networks regained an unbiased, symmetrical representation of both outcomes (symmetric “x” shape), while lazy networks remained unaffected.

The computational principle outlined above suggests a working hypothesis for the function of paradoxical replay: it serves to protect rich, but not lazy, task representations from destructive interference due to unbalanced training. This hypothesis predicts that on tasks or situations characterized by low-dimensional, context-dependent “rich” representations, paradoxical replay should be strong because it serves an important adaptive function. Conversely, on tasks that favor high-dimensional, context-independent “lazy” representations, there is less need for paradoxical replay and it will be weak. To test this idea, we first quantify the representational similarity of hidden unit activity in rich vs. lazy networks. Using both a single-cell measure, choice selectivity for each individual unit (Figure 3a) and an ensemble level measure, populational representational distance (Figure 3c), we found a stronger separation between hidden activity encoding the two choices in rich networks compared to lazy networks (Figure 3b, d). These differences did not exist in rich networks upon initialization, indicating their development during training rather than being inherent in the initial network weights (Figure S3). The choice-dependent activity of rich network hidden units is analogous to the widely studied “splitter cells” and their ensemble counterparts in the rodent hippocampus (Ferbinteanu & Shapiro, 2003; Wood et al., 2000; see Duvelle et al., 2023 for review).

**Figure 3:**
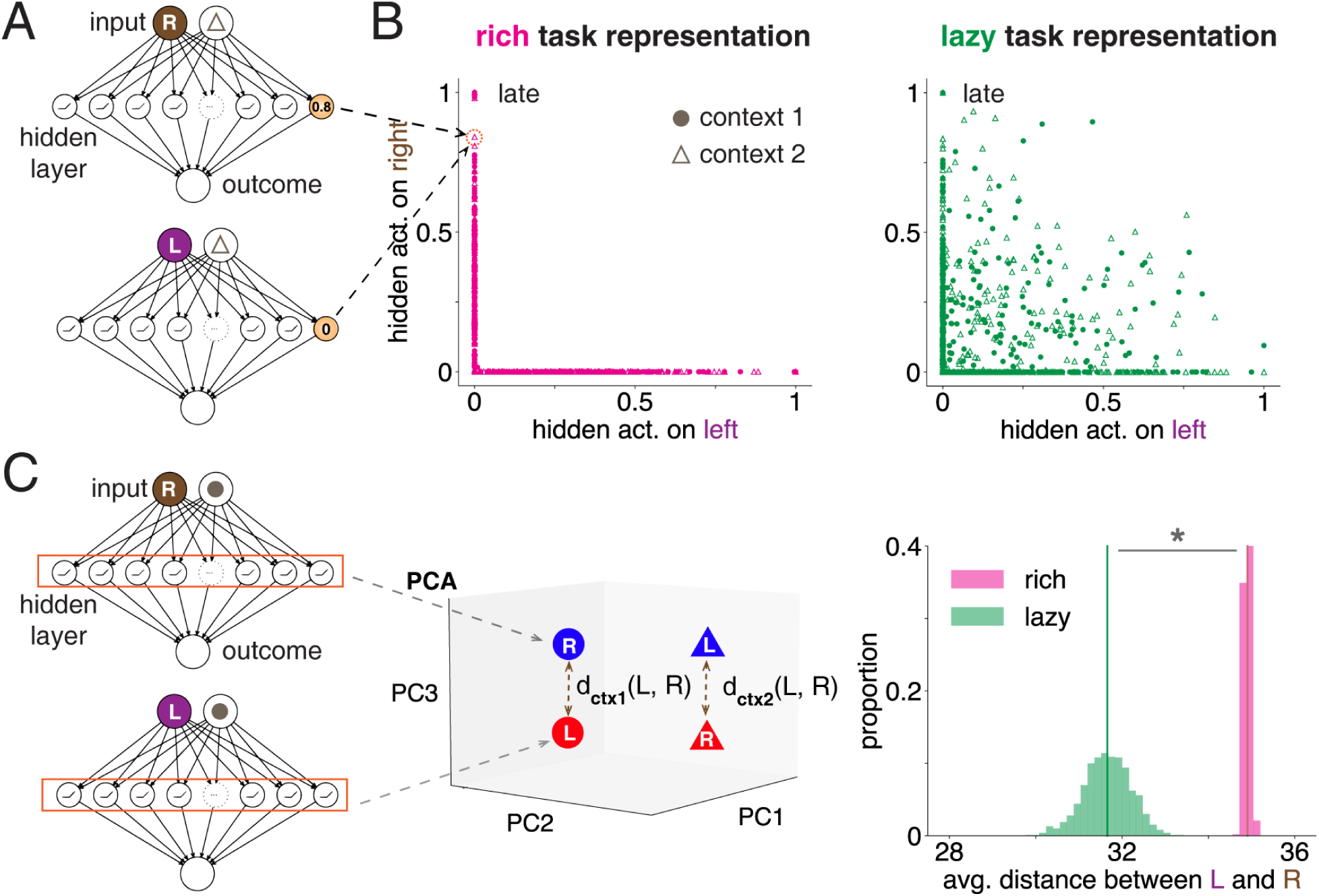
Emergence of context-dependent “splitter” representations in rich but not lazy networks. **(A)** Schematic of example unit activation in the hidden layer of the network for both left and right choices. **(B)** Rich networks exhibit selective activation (“splitting”) for either left or right choices, while lazy networks have mixed encoding of both choices. **(C)** Illustration showing a 3D representation of populational activations in the hidden layer for left and right choices under each context. **(D)** Rich networks demonstrate larger distances between left and right (averaged across contexts) compared to lazy networks (p < 0.001, Wilcoxon rank-sum test), indicating stronger splitter strength in the ensemble hidden unit activity pattern.

Next, we applied these measures of rich task representations to data recorded from the rodent hippocampus using the Carey et al. (2019) data set (see Figure 4a for illustration and Figure S1a for task schematic). Specifically, for each individual recording session, we quantified the “richness” of the active task representation using both the single-cell splitter strength (averaged across cells) and the ensemble representational distance by comparing neural activity for left and right trials. This representational richness measure was then correlated with the strength of paradoxical replay (see Figure 4b for a schematic of how we quantified this). The clear prediction from our theory is that when a highly contextual, “rich” representation (high splitter strength) is in effect, paradoxical replay will be strong; conversely, more task-agnostic, “lazy” representations (low splitter strength) should be associated with orthodox replay. Strikingly, those sessions in which rats showed the largest representational separation showed the strongest paradoxical replay (Figure 4c, d), both for single-cell (r = 0.53, p < 0.05) and ensemble (r = 0.49, p < 0.05) measures of representational similarity. Importantly, we verified that this effect also held true when paradoxical replay and richness of representation were estimated on non-overlapping areas of the maze to avoid circularity in the analysis (single-cell splitter: r = 0.50, p < 0.05; ensemble splitter: r = 0.46, p < 0.05), and when only significance-thresholded replay events were considered (single-cell splitting: r = 0.55, p < 0.05; although the positive correlation for ensemble splitting did not reach statistical significance: r = 0.32, p = 0.18). Note that comparisons between left and right trajectories were made using equal numbers of trials to avoid biases in our ability to detect replay due to unbalanced experience (van der Meer et al. 2020; see *Methods* for details). To further test the robustness of this result, we considered different normalized measures of splitting beyond the raw firing rate difference between left and right trials used in the main analysis; although the significance level depended on the normalization used (firing rate difference divided by sum: r = 0.41, p = 0.083; divided by euclidean distance (L2 norm): r = 0.57, p = 0.011) the direction and magnitude of the correlation remained consistent.

**Figure 4:**
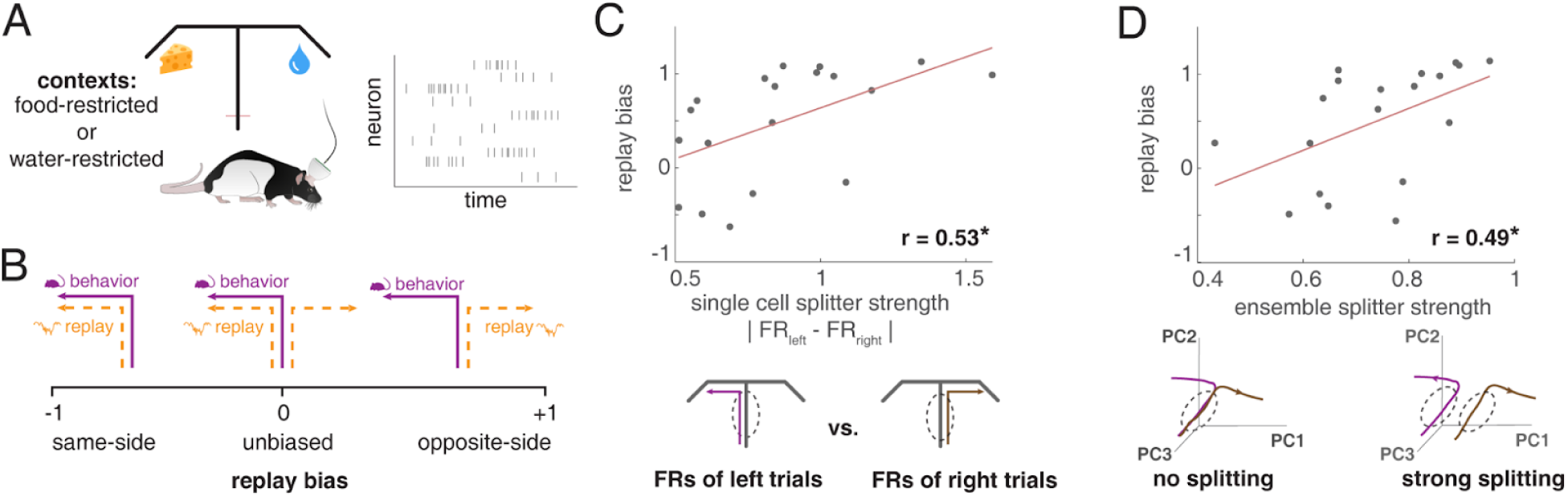
Paradoxical replay correlates with both single-cell and population measures of rich representations, as quantified by “splitter” strength. **(A)** Illustration of a rat performing a T-maze task in Carey et al. (2019), with implanted tetrodes in hippocampal CA1. Sessions alternated daily between food-restriction and water-restriction, causing unbalanced experience. **(B)** Schematic of how we quantified paradoxical replay (see *Methods* for details): a higher score indicating replay *opposite* to behavioral choice in a given session. **(C)** Sessions with larger single-cell representational distance (defined by absolute difference in firing rates) indicative of rich representations showed stronger paradoxical replay (r = 0.53, p < 0.05). **(D)** Paradoxical replay bias also positively correlated with greater separation between left and right ensemble firing patterns (r = 0.49, p < 0.05).

Next, we sought to test the hypothesized link between splitter strength and replay content in an independent data set. Gupta et al. (2010) examined replay content as rats navigated a continuous T-maze with reward contingencies that switched once within each recording session (between “left rewarded”, “right rewarded” and “alternation rewarded” blocks). This experimental design provides a strong test of our hypothesis because, in those recording sessions that included alternation, there is a within-session comparison of balanced experience (alternation) and unbalanced experience (left, or right). The key result from the Gupta et al. study is that during left (or right) blocks, rats replay the opposite (non-chosen) side of the maze more than during alternation blocks. This is surprising because during alternation, the opposite side has been visited more recently (and will be visited again sooner) compared to left (or right) blocks, demonstrating that on that task, replay does not track recent or upcoming choices. Mapping this result onto our working hypothesis, the prediction is that those blocks with more paradoxical replay (i.e. the left or right-only blocks) have a richer task representation, as measured by stronger splitter strength, compared to alternation blocks.

In accordance with this prediction, left (L) or right (R) blocks showed significantly higher splitter strength compared to alternation (Alt) blocks (Figure 5, left panel; p < 0.001, Wilcoxon). Replotting the data from Gupta et al. (2010, Figure 5) shows the match of this novel splitter strength difference with the previously reported paradoxical replay difference (i.e. more replay of the opposite, non-chosen side during L or R; Figure 5, right panel). As before, we further tested the robustness of this effect by applying different normalization methods, and found the results to be consistent despite some variation in significance levels (firing rate difference divided by sum: Alt = 0.362 ± 0.030, R/L = 0.460 ± 0.041, p = 0.084, euclidean distance: 0.368 ± 0.041, R/L = 0.649 ± 0.070, p = 0.003). Thus, despite substantial differences between the Carey et al. and Gupta et al. tasks (e.g. continuous vs. discrete trial maze; motivational shifts vs. task contingency switch; on-task vs. off-task replay), we found evidence for the a *priori* predicted link between splitter cell strength and opposite-side replay in both.

**Figure 5:**
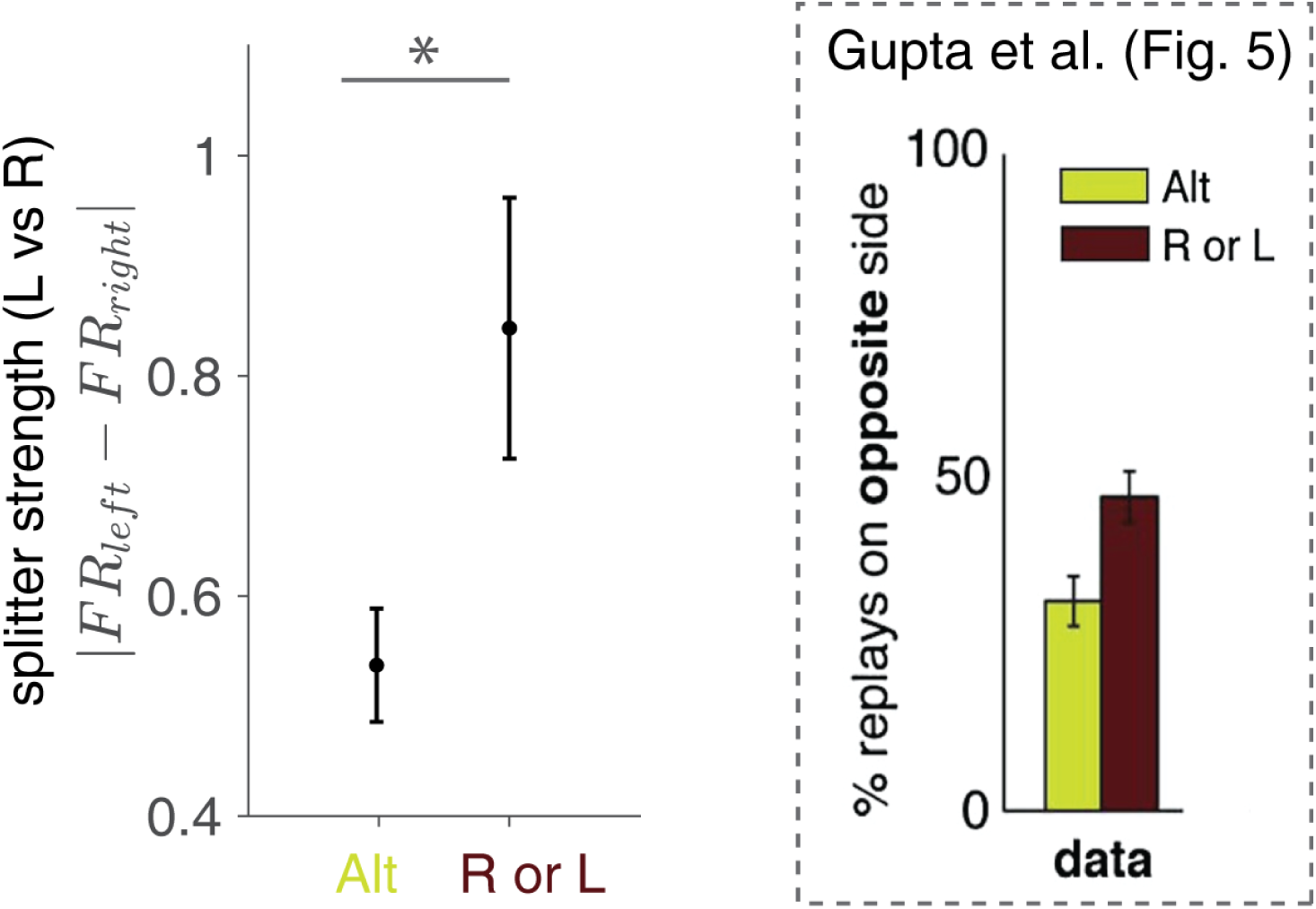
Splitter strength correlates with paradoxical replay in the Gupta et al. dataset. (**Left column**) Splitter strength, indicated by the differential firing rates of individual cells during left (L) vs. right (R) running trajectories, is significantly lower during trial blocks that require alternating (Alt) between L and R compared to repeating the same trajectory. (**Right column**) Paradoxical replay is more prevalent in blocks where repeating the same L or R trajectory was rewarded, compared to blocks where alternating between L and R was rewarded (adapted from Gupta et al., 2010). Thus, higher splitter strength and stronger paradoxical replay co-occur in this study.

## Discussion

In this study, we provide a normative theory to explain the phenomenon of “paradoxical replay”, supported by proof-of-principle simulations that reveal a computational tradeoff and that generate novel predictions, which we then tested and confirmed with new analyses of experimental data from two different studies. Specifically, we first demonstrate that in a toy computational model, the usefulness of paradoxical replay in protecting memories from destructive interference depends on (1) the degree of unbalancedness of experience, and (2) the nature of the task representation in use: “rich” but not “lazy” task representations are susceptible to detrimental effects of repeated, stereotyped experience. This principle predicts that paradoxical replay in the brain should occur preferentially when rich representations meet unbalanced experience, a prediction we confirm experimentally by demonstrating a positive correlation between the paradoxicality of replay and neural signatures of rich representations in the rodent hippocampus.

Most immediately, our results address an outstanding puzzle in the literature. Multiple findings spanning rats (Carey et al., 2019; Gillespie et al., 2021; Gupta et al., 2010) and humans (Wimmer et al., 2023) have demonstrated paradoxical dissociations between behavioral preference (experience) and replay content. For instance, in Carey et al. rats consistently replayed the opposite maze arm from the one they behaviorally preferred, and in Gillespie et al. (among other results) the previous, no-longer rewarded arm was replayed more. These results are inconsistent with the leading theories of replay content and function, including various flavors of the “two-stage” memory model in which hippocampus serves as a time-limited memory buffer for systems consolidation (Buzsáki, 1989; Kumaran et al., 2016; McClelland et al., 1995); proposals that replay implements a decision-making or planning process in service of upcoming behavior (Eldar et al., 2020; Ólafsdóttir et al., 2017; Pfeiffer & Foster, 2013) and the idea that replay implements “fictive learning” that fixes those reward prediction errors in value maps that are most useful for future behavior (Mattar & Daw, 2018; Sagiv et al., 2025; Sutton, 1991).

Although none of these canonical replay theories account for paradoxical replay, two recent studies have explored this issue with state-of-the-art modeling approaches that cover an impressive range of experimental results. Sagiv et al. (2025) extended the Mattar & Daw (2018) approach to situations where animals learn the value functions of multiple different maps, one map for each goal location. If the animals learn about these goals (across maps) at a different rate than they learn about values within maps, then the two processes can become desynchronized and paradoxical replay can appear as a side effect of switching goals at a specific rate. Zhou et al. (2026) conceptualize replay as probabilistically jumping to contextually related experiences, enabling offline association and inference. Paradoxical replay occurs in this model by adding a mechanism whereby online experience can inhibit replay of that experience (see Mallory et al. 2025), but this leaves open how such inhibition would be useful. Thus, while these recent models do account for a wealth of experimental data on the content and function of replay, paradoxical replay appears as a side effect rather than as a phenomenon with a specific adaptive function. In contrast, the current work offers a different, normative perspective on the function of paradoxical replay, rooted in an understanding of the structure, at the neural coding level, of task representations -- a connection that has been missing from theories of replay to date.

In this study we tested predictions from our proposed function of paradoxical replay by taking advantage of within-session changes in experience sampling (Gupta et al. 2010), or between-session and between-subject variability (Carey et al., 2019). Importantly though, we believe that our theory and its predictions fit the pattern of replay content across many studies. In particular, those studies where paradoxical replay has been most clearly seen have in common that the animal performs blocks of repeated, stereotyped trials (Carey et al., 2019; Gillespie et al., 2021; Gupta et al., 2010; see Table 1). These are precisely the conditions conducive to rich, contextual task representations as shown by the prevalence and strength of “splitter cells” that distinguish the blocks during overlapping segments (Duvelle et al., 2023; Ferbinteanu & Shapiro, 2003; Wood et al., 2000). In contrast, studies that have not reported paradoxical replay tend not to switch between different blocks but instead interleave different trials (e.g. T-maze alternation) or encourage learning of a single, non-contextualized map by having the animal run all trajectories in all directions, as occurs in e.g. open fields (e.g. Pfeiffer & Foster 2013). Thus, the principle revealed by our model fits the broader pattern of experimental results in the literature, and can be used to make novel predictions. For instance, in the commonly used W-track alternation task (Frank et al. 2000; Singer et al. 2013; Shin et al. 2019) animals alternate between the left and right arms on outbound trials but return to the center arm on inbound trials, resulting in both balanced (outbound; approximately equal left and right trials) and unbalanced (inbound, usually center arm but sometimes outer arm) experience regimes. Our account predicts that on inbound trials (e.g. left > center) replay of the opposite arm (right) is more likely than on outbound (center > left) trials, an idea that can be tested experimentally.

Our model has a number of limitations. A specific open issue is that we modeled paradoxical replay as “re-balancing” unbalanced experience. While this account explains some features of the Gillespie data, it does not match the observed bias to the previously rewarded goal, or the gradual increase of replay of the rewarded arm. Thus, rather than rigidly replaying all possible but non-chosen options, a more nuanced definition of paradoxical replay may be to replay representations whose behavioral sampling is currently insufficient relative to their learned importance or expected future usefulness. Indeed, in order to keep a clear focus on the function of paradoxical replay and the situations it would be predicted to occur, our model does not attempt to incorporate known factors that shape the content of replay, such as the influence of reward and novelty (Cheng & Frank, 2008; Dupret et al., 2010; Ambrose et al., 2016; Michon et al., 2019) -- which Sagiv et al. (2025), Zhou et al. (2026) and related work (Mattar & Daw, 2018; Momennejad et al., 2018; Jensen et al. 2024) does incorporate, although as discussed above even these accounts do not provide a fully settled explanation. In common with Sagiv et al., we do not provide a mechanistic explanation for how the hippocampus would “know” what to replay; however, the phenomenon of “rebound” in SWR events after closed-loop suppression in the hippocampus (Gillespie et al., 2024; Girardeau et al., 2014) shows that there is a SWR pressure that is under homeostatic control. Further experimental work can determine what the physiological basis of this pressure is, whether it is content-specific, and inform how it is best modeled.

More generally, by considering the different ways that artificial and biological networks can represent and solve model tasks, with rich and lazy learning as paradigmatic examples, we demonstrate that the effects of replay interact with the nature of task representations. As such, our work has implications beyond merely addressing a puzzle in rodent replay data: we refine the commonly used idea of replay as a mechanism to prevent catastrophic interference in “continual learning” and related scenarios (McClelland et al., 1995; van de Ven et al., 2020) to be more nuanced and dependent on experience and on task representations. Furthermore, this novel link between replay and task representations suggests possible predictions and avenues for further work: for instance, the intriguing phenomenon of representational drift (Driscoll et al., 2022; Rule et al., 2019; Ziv et al., 2013), would be expected to be larger under lazy learning conditions compared to rich learning, and therefore should correlate inversely with paradoxical replay. Big-picture, a complete understanding of the likely varied roles of replay in the brain will require bridging different literatures and perspectives, an objective the present study contributes to by linking the format of task representations to replay.

## Methods

### RESOURCE AVAILABILITY

#### Lead contact

Further information and requests for resources and reagents should be directed to and will be fulfilled by the lead contact, Matthijs A. A. van der Meer.

#### Materials availability

This study did not generate new unique reagents.

#### Data and Code Availability

The Carey dataset used in this study is publicly available from DataLad (http://datasets.datalad.org/?dir=/workshops/mind-2017/MotivationalT). Code necessary and sufficient for reproducing all results and figures in this manuscript is publicly available on Github (https://github.com/vandermeerlab/replay_task_rep).

### EXPERIMENTAL MODEL AND SUBJECT DETAILS

In addition to neural network simulation and analysis, this study used experimental data from two previously published data sets. The Carey et al. (2019) study consisted of 4 male Long–Evans rats, aged 4–8 months at the start of behavioral training, and then chronically implanted with arrays of recording electrodes. Gupta et al. (2010) contained 6 male Fisher-Brown-Norway hybrid rats, aged 7–10 months at time of implantation.

### METHOD DETAILS

#### Neural network simulations

We performed and analyzed neural network simulations using Python, employing the NumPy, SciPy, PyTorch, Scikit-Learn and Pingouin packages.

##### Overall approach and task design

We modeled a canonical contextual discrimination task where the outcomes of choices such as left (L) or right (R) reversed between two contexts (context 1: L+R-, context 2: L-R+; Figure 1a). The network was trained to predict outcomes {+,-} based on choices {L,R} in each context. We built networks initialized with different input-to-hidden weights and trained with different proportions of input patterns and compared the resulting weight and hidden unit activation patterns, all described in more detail below.

##### Neural network architecture

Specifically, we used a 3-layer feedforward neural network architecture consisting of input, hidden, and output layers. Inputs included choice (left or right) and context (context 1 or context 2). These inputs were mapped to a hidden layer of 500 units, passing through Rectified Linear Unit (ReLU) nonlinearities. ReLU outputs projected onto a linear output unit that predicted outcome valence (−1 or +1).

##### Weight initialization (rich vs. lazy learning)

We initialized all network parameters with random sampling from Gaussian distributions (mean zero). To induce learning of rich vs. lazy representations, we varied the standard deviation of input-to-hidden weights (0.0025 for *rich* and 0.25 for *lazy*). Hidden-to-output weights in both regimes had a standard deviation of 1 / **N_hidden_**, where **N_hidden_** is the number of hidden units.

##### Training

We conducted 20 independent runs for both rich and lazy learning regimes, training the network with gradient descent (learning rate 0.002, SGD optimizer in PyTorch) over 5000 iterations. Models were trained on Mean Squared Error (MSE-Loss) between true and predicted outcomes.

##### Balanced vs. unbalanced experience

To simulate the effects of balanced vs. unbalanced experience, we changed the distribution of inputs the networks were trained on. In the balanced distribution (Figure 1) all choice-context combinations occurred with equal frequency, while in the unbalanced distribution (Figure 2) positive outcomes were four times more frequent than negative outcomes (4+1-), similar to the frequencies empirically observed in Carey et al. (2019).

##### Paradoxical replay simulation

To simulate paradoxical replay, we provided networks trained with unbalanced experience, as described above, with additional training data of the less-experienced condition (1+4-). Learning rate was the same for actual vs. replayed experience.

##### Addition of Gaussian input noise

We investigated the robustness of rich vs. lazy representations by adding Gaussian noise to the inputs (Figure S1c). Twenty independent runs for both regimes were conducted with Gaussian noise added to input units at the testing stage (once networks had converged) with the SD varied parametrically in 20 steps from 0.1 to 2.

#### Rat behavioral task

We performed new analyses on the Carey et al. (2019) and Gupta et al. (2010) datasets, comprising ensemble recordings of hippocampal CA1 neurons from rats navigating T-maze tasks. Rats chose between the left and right arms of the maze during daily sessions. In Carey et al., rats experienced alternating food and water restrictions across days: the left arm offered a food reward (five 45 mg pellets), while the right arm provided a water reward (∼0.2 ml sucrose solution). Each recording session consisted of 15-20 discrete trials, with at least 5 trials for the less preferred choice (left or right). In Gupta et al., daily sessions used a continuous “two-T” maze where rats were rewarded according to one of three contingencies: left-only, right-only, or alternation at the final choice point. During training, the reward contingency was stable within a session; during recording sessions, the contingency switched approximately midway through the session.

#### Hippocampal neural data analysis

Neural data analyses were conducted in MATLAB 2022b (Mathworks).

##### Criteria for inclusion of data

As in our previous work (Carey et al., 2019; Chen et al., 2021; van der Meer et al., 2017) only sessions with at least 40 simultaneously recorded neurons were included, leaving 19 out of 24 total sessions for analysis. Putative interneurons (mean firing rate > 5 Hz) were excluded from the analysis.

##### Splitter signal

To assess how left and right choices were separately represented, we compared firing rates between left and right trials using data *only from the shared central stem of the maze*.

##### SWR detection and replay content decoding

Sharp-wave ripple (SWR) events were detected based on ripple LFP power and multi-unit activity as described in Carey et al. (2019). Events that exceeded the detection threshold were identified as potential replay events. Next, neural activity within each event was decoded as a single time window using a standard memoryless Bayesian decoding approach. Spatial tuning curves used for decoding were estimated from running data either *across the entire maze* or specifically on the *maze arm past the choice point*, with equal numbers of left and right trials. Full details of the detection and decoding procedures were outlined in Carey et al. (2019). For the Gupta et al. (2010) analysis, we reproduced the relevant figure from the original paper (Figure 5b).

### QUANTIFICATION AND STATISTICAL ANALYSIS

#### Neural network simulations

##### Representational similarity analysis (RSA) of hidden unit activity

To evaluate how left and right choices were differently represented in rich vs. lazy networks during training, we conducted RSA on hidden layer activity patterns. For each run, we performed principal component analysis (PCA) with three principal components to project hidden unit activations for each choice-context combination into an embedding space. Subsequently, we computed the Euclidean distances between left and right activations, averaged across contexts, and compared results between rich and lazy regimes across 1000 runs. A larger distance in the embedding space indicates that left and right choices are represented more separately.

#### Hippocampal neural data analysis

##### Carey et al. single-cell and ensemble splitter strength

Splitter strength was defined as differential firing between left and right trials along the common central stem of the T-maze. Specifically, single-cell splitter strength for a given session was quantified as the absolute difference in firing rates (FR), averaged across neurons, while population splitter strength was calculated through classification accuracy between left and right choices using a Bayesian decoder. Higher values in both metrics indicate more differential firing patterns and thus greater splitting.

##### Carey et al. paradoxical replay bias

Each SWR event, detected and decoded as described above, was categorized as representing the *opposite* or *same* as the preferred behavioral choice in that session. This categorization was based on a z-score comparing the decoded probability to a shuffled distribution generated from 1000 shuffles, where the left and right tuning curves of individual neurons were randomly permuted. Two measures were computed: 1) the average z-score bias across *all* replay events and 2) the proportion of events exhibiting opposite-side bias out of *significance-thresholded events*. In both cases, a higher score suggests a greater bias in replay events relative to the preferred behavioral choice in that session. For example, replay predominantly represented the right choice when rats primarily chose the left arm, as in a food-restricted session.

##### Gupta et al. splitter strength

Each session was divided at the contingency switch, and task blocks with more than 10 trials were retained. Single-cell splitter strength was computed separately for each retained task block as the absolute difference between mean firing rates on left and right trajectories, averaged across neurons. For repeat blocks, splitter strength was included only when more than two error trials were available, allowing left-right firing-rate differences to be estimated under the repeat contingency. Splitter strength was then compared between alternation blocks and repeating blocks.

##### Gupta et al. paradoxical replay bias

We reproduced the published Gupta et al. result, in which opposite-side replay was more prevalent during repeat blocks than alternation blocks.

## Acknowledgements

We thank Jeremy Manning, John Murray, James Fitzgerald, Kari Hoffman, and Cristina Savin for their helpful feedback and discussion. This work was supported by NIMH R01 MH123466 to MvdM.

## Author contributions

H.-T.C. and M.A.A.v.d.M. conceived the project, brainstormed analysis ideas, and discussed the results. H.-T.C. implemented neural network simulations and all analyses. H.-T.C. and M.A.A.v.d.M. jointly wrote the paper.

## Supplementary Information

**Figure S1:**
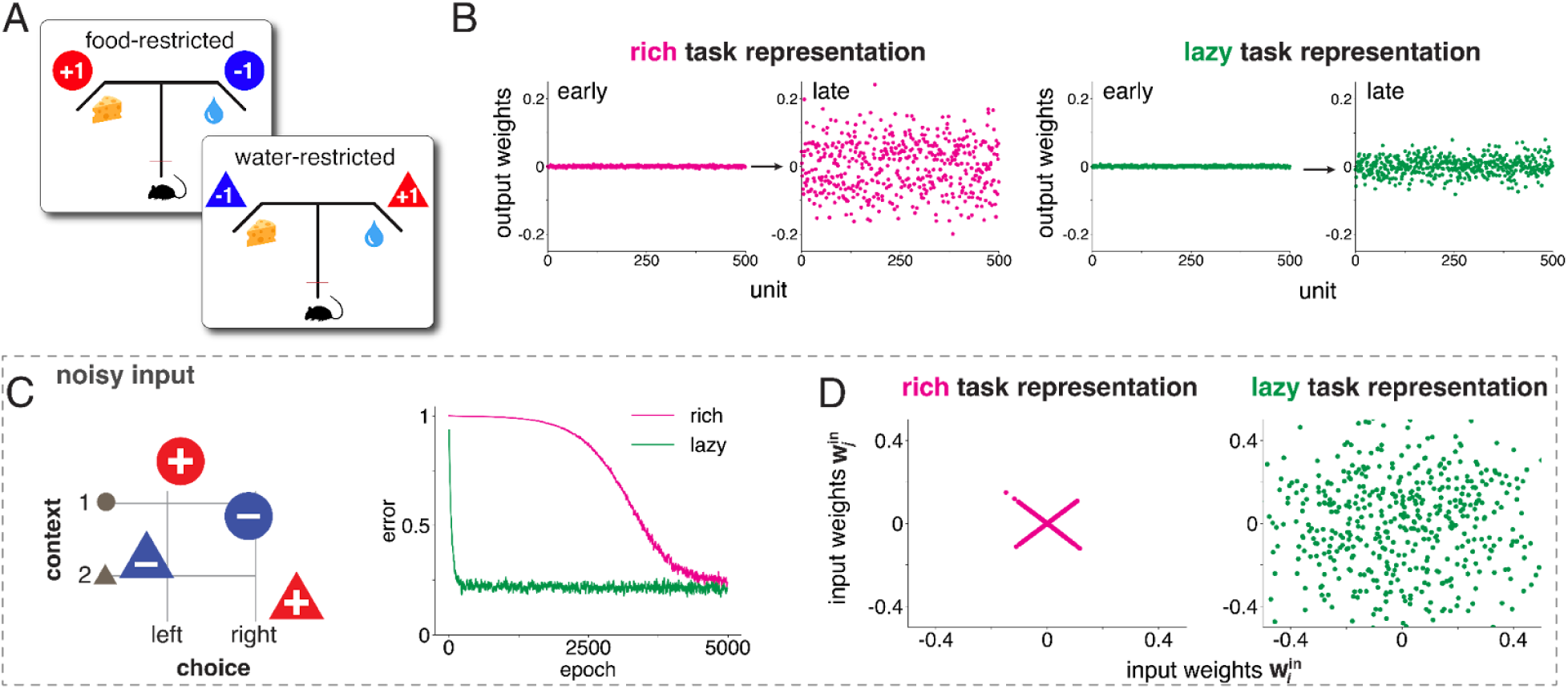
Carey task schematic and trade-off between robustness and adaptability in rich vs. lazy learning. **(A)** In Carey et al. (2019), rats underwent alternating water and food restriction prior to daily recording sessions. During food-restricted sessions, rats preferred the arm with a food reward (*left, +1*) in a T-maze while avoiding the arm with a water reward (*right, −1*). Conversely, during water-restricted sessions, these preferences reversed (*left, −1* and *right, +1*). **(B)** Hidden-to-output weights in rich networks underwent more substantial changes during training compared to lazy networks. **(C)** Rich-learning and lazy-learning networks were trained in a noisy setting, where inputs were randomly sampled in each training epoch (example inputs shown in the left panel). Lazy-learning networks exhibited faster learning compared to rich-learning networks. **(D)** Representations in the input-to-hidden weights closely resemble those trained with deterministic inputs (Figure 1e, f, right panel).

**Figure S2:**
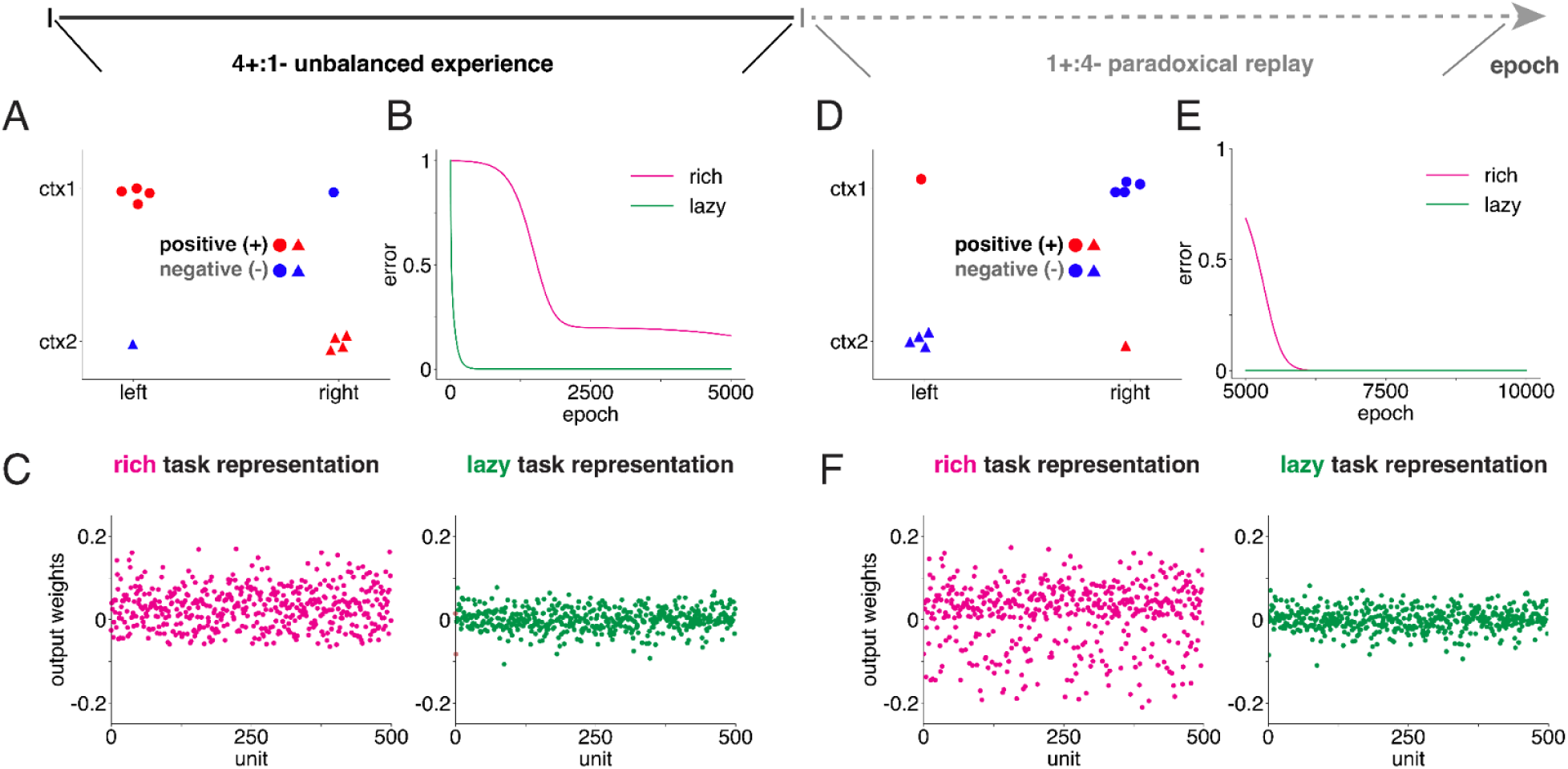
Implementation of unbalanced training in neural network models. **(A)** We implemented biased training by adding disproportionate training samples of positive outcomes. **(B)** Networks initialized in the lazy regime showed faster adaptability in handling unbalanced data distribution, while rich-learning networks failed to cope with this situation within the limited training time. **(C)** Output weights in rich-learning networks exhibited a bias toward positive values due to the overrepresentation of positive outcomes (see Figure S1d for comparison with balanced input training). Conversely, output weights in lazy-learning networks remained unbiased. **(D)** Biased replay of the negative contingency was implemented by changing the majority class from positive outcomes to negative outcomes after biased training of the positive contingency. **(E)** Introducing biased replay helped rich-learning networks in learning the task and achieving performance comparable to lazy-learning networks. **(F)** Output weights in rich-learning networks became unbiased after introducing paradoxical replay bias.

**Figure S3:**
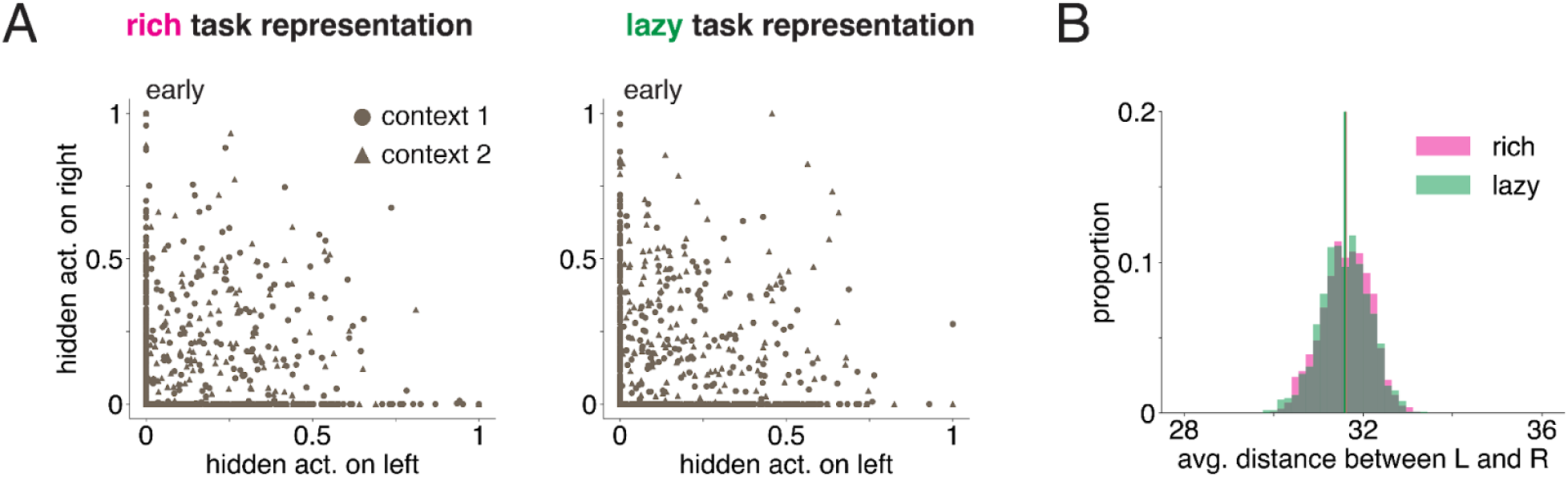
Initial representations in hidden-layer activations are similar across rich and lazy networks. **(A)** During the early stage of training, hidden-layer units in both rich- and lazy-initialized networks exhibited a comparable degree of non-selective activation for both left and right choices. **(B)** Left and right choices are similarly represented in the population activations of hidden layers between rich-learning networks and lazy-learning networks. (p = 0.29 for Wilcoxon rank-sum test).

